# Histone H1.2 Dependent Translocation of Poly (ADP-ribose) Initiates Parthanatos

**DOI:** 10.1101/2023.06.18.545460

**Authors:** Jing Fan, Bong Gu Kang, Tae-In Kam, Adam A. Behensky, Jesse Rines, Ho Chul Kang, Valina L. Dawson, Ted M. Dawson

**Author notes:** Authors contributed equally (Valina L. Dawson, Ted M. Dawson). Authors contributed equally (Jing Fan, Bong Gu Kang, Tae-In Kam). Correspondence to: Ted M. Dawson, M.D Ph.D. or Valina L. Dawson, Ph.D.

## Abstract

Toxic cellular insults activate the nuclear protein poly (ADP-ribose) (PAR) polymerase-1 (PARP-1) to initiate parthanatos, a regulated cell death program. PAR acts as a death signal by translocating from the nucleus to the cytosol, where it activates the next steps in the parthanatic cell death cascade. How PAR translocates from the nucleus to the cytosol is not known. Here we show that PARylation and PAR binding to histone H1.2 enables it to act as a carrier, transporting PAR out of the nucleus to the cytosol. Knocking down the expression of histone H1.2 via CRISPR/Cas9 and knockout of histone H1.2 reduces the translocation of PAR to the cytosol after treatment of human cortical neurons with N-methyl-D-aspartate (NMDA) or following oxygen-glucose deprivation (OGD). The PAR-dependent E3 ubiquitin ligase, Iduna (RNF146) ubiquitinates PARylated H1.2. Overexpression of Iduna reduces the expression levels of cytosolic histone H1.2, preventing the translocation of PAR following NMDA or OGD exposure, similar to inhibition of PAR formation by the PARP inhibitor, DPQ. Whereas, the catalytically null variant Iduna C60A, or the PAR binding mutant Iduna Y156A and R157A (YRAA) was ineffective in ubiquitinating histone H1.2 and preventing the reduction in cytosolic histone H1.2 levels and PAR translocation from the nucleus to the cytosol. Histone H1.2 heterozygote and homozygote knockout mice exhibited reduced infarct volume 24 hrs post middle cerebral artery occlusion (MCAO) and showed better recovery in motor deficits than wildtype littermates at day 3 and/or day 7 post MCAO. Collectively, these findings reveal histone H1.2 as the key carrier of PAR out of the nucleus to the cytosol where it participates in the next step of the parthanatic cell death cascade.

## Introduction

Parthanatos is a regulated cell death pathway that is due to excessive activation of poly (ADP-ribose) (PAR) polymerase-1 (PARP-1) ^1–4^. Inactivation or inhibition of PARP is dramatically protective in a variety of cell injury paradigms, including glutamate excitotoxicity, stroke, ischemia-reperfusion injury, Parkinson’s disease (PD) among others ^1–6^. PARP1 functions in the DNA damage response to regulate DNA repair by ribosylating proteins and acting as a scaffold for the cellular machinery that repairs genomic DNA ^7–10^. When over activated PARP-1 generates large-branched chain (PAR) polymers that are toxic ^11^. These toxic PAR polymers are generated in the nucleus but exit to act outside the mitochondria to inhibit mitochondrial hexokinase ^2, 12, 13^ and release apoptosis inducing factor (AIF) from the mitochondria ^14–16^. Inhibition of hexokinase by PAR leads to energy depletion via inhibition of glycolysis ^12, 13^. AIF translocates to the nucleus and carries the parthanatos associated AIF nuclease (PAAN), macrophage inhibitory factor (MIF) to initiate large scale DNA fragmentation, which is the terminal event in parthanatos ^17–19^.

Some studies suggest that free PAR exits the nucleus via the cleavage of PAR from PARylated proteins by PAR glycohydrolase (PARG) and ADP-ribosyl-acceptor hydrolase 3 (ARH3) ^20^. How PAR, which is a large negatively charged molecule, could exit the nucleus to mediate parthanatos has been a mystery. It is likely that there is a ribosylated or PAR binding protein that facilitates the transfer of nuclear PAR to the cytoplasm to initiate parthanatos.

There are many nuclear proteins that are PARylated or bind PAR itself that could be candidates for carrying PAR out of the nucleus ^21–31^. One of the major classes of proteins that are PARylated and bind PAR are histones ^23, 26, 32^. Of the histones, the most abundant that are extensively ribosylated are the histone H1 family members ^33^. Ribosylation of histone H1 relaxes the chromatin structure to facilitate DNA repair ^34–36^. There are at least five histone H1 isotypes ^37^. Among the histone H1 isotypes, histone H1.2 exits the nucleus in response to DNA double-strand breaks to initiate cell death ^38, 39^. Here we show that histone H1.2 carries PAR out of the nucleus to the mitochondria where it participates in parthanatos.

## Results

To determine if histone H1.2 binds PAR, a PAR overlay assay and a PAR binding gel shift assay were performed and showed that histone H1.2 robustly binds PAR *in vitro* (Fig. S1A). Histone H1, H1.3, H2A, H2B, H3 and H4 also bind PAR (Fig. S1A). Histone H1.2 is also ribosylated by PARP-1 as determined by an *in vitro* ribosylation assay (Fig. S1B). Previously we showed that human cortical neurons die primarily by parthanatos following glutamate excitotoxicity or oxygen-glucose deprivation (OGD) ^40^. In this study, we observed that one hr after stimulation of human cortical neurons by N-methyl-D-aspartate (NMDA) or OGD, immunoprecipitation of PAR pulls down histone H1.2 and immunoprecipitation of histone H1.2 pulls down PAR indicating that histone H1.2 is either ribosylated or binds PAR following neurotoxic NMDA or OGD exposure (Fig. 1A-B). To monitor whether histone H1.2 and PAR translocate out of the nucleus, human cortical neurons were infected with lenti-GFP to demarcate the cytosol and exposed to neurotoxic concentration of NMDA or OGD. The cultures were assessed by immunohistochemistry at 15 minutes or one hr following exposure. Both PAR and histone H1.2 are visualized in the cytosol of both NMDA (Fig. 1C-D) and OGD (Fig. 1E-F) treated human cortical cultures within 15 minutes, and both remain in the cytosol for up to one hr after exposure to NMDA or OGD. 3,4-Dihydro-5-[4-(1-piperidinyl)butoxyl]-1(2H)-isoquinolinone (DPQ), a PARP inhibitor, completely prevents the formation of PAR and the translocation of histone H1.2 to the cytosol (Fig. 1C-F). These observations were confirmed by subcellular fractionation and immunoblot analysis (Fig. 1G-J).

**Fig. 1.**
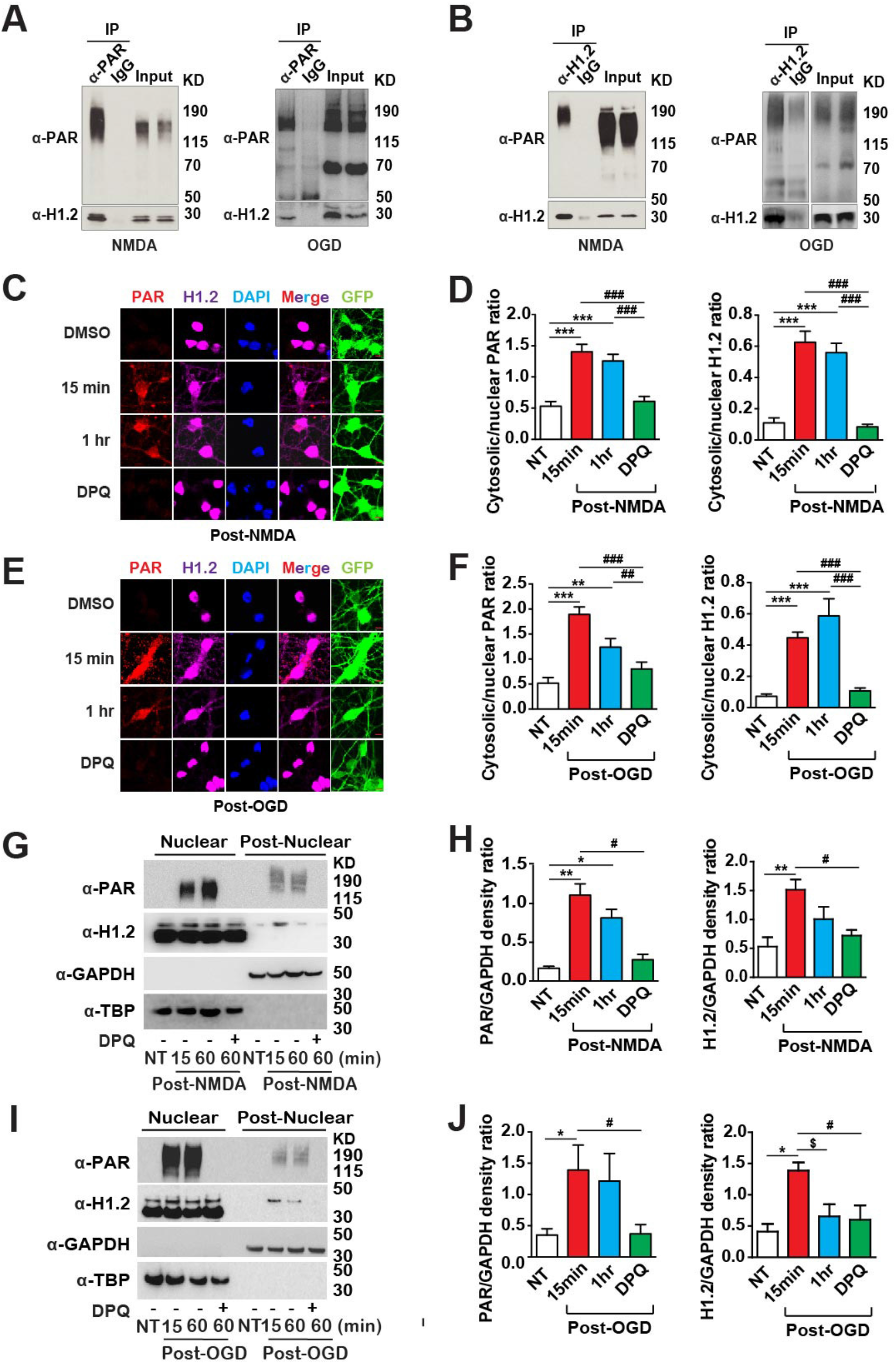
Histone H1.2 interacts and co-translocates with PAR from nuclei to cytosol/mitochondria after NMDA and OGD stimuli. (**A, B**) Representative immunoblots showing H1.2 interacts with PAR or poly-ADP ribosylated proteins in human cortical neurons after NMDA or OGD stimulation. Human neuron lysates are incubated with (**A**) anti-PAR or (**B**) anti-H1.2 antibodies, and rabbit or mouse IgG respectively for immunoprecipitation. Blots are cut along 50KD, and probed with anti-PAR antibody and anti-H1.2 antibody respectively. (**C-F**) Representative photomicrographs showing human cortical neurons transduced with lentiviruses carrying GFP. Cells were fixed with 4% PFA for 15 minutes and 1 hr after NMDA (**C**) or OGD (**E**) challenge with or without DMSO or DPQ treatment, stained with anti-PAR antibody (red), anti-H1.2 antibody (magenta) and DAPI (blue). The merged panel is a composite of the red, magenta and blue channels. The cytosolic and nuclear levels of H1.2 and PAR in (**C**) and (**E**) are quantified and the ratios (translocation) are expressed as mean ± s.e.m of 15 independent cells from 3 independent experiments in (**D**) and (**F**), respectively. *P*<0.001 by one-way ANOVA for both PAR and H1.2 translocations in NMDA (**C**) and OGD (**E**) experiments, \*\**p*<0.01, \*\*\**p*<0.001, ^##^*p*<0.01, ^###^*p*<0.001 by Bonferroni’s posttest between groups indicated on the bar graph. Scale bar = 10 µm. (**G-J**) Representative immunoblots showing human cortical neurons lysates harvested before, or 15 minutes, or 1 hr after NMDA (**G**) and OGD (**I**) challenge with or without DPQ, and subcellular fractionated. Blots were probed with anti-PAR antibody, anti-H1.2 antibody and markers for nuclear and cytosolic fraction markers (TBP and GAPDH). NT, no treatment. The post-nuclear levels of H1.2 and PAR are quantified and the ratios to GAPDH (translocation) are expressed as mean ± s.e.m of 4-7 independent experiments in (**H**) and (**J**). *P*<0.001 by one-way ANOVA for PAR and H1.2 translocations in NMDA (**G**) and OGD (**I**) experiments, \**p*<0.05, \*\**p*<0.01, ^$^*p*<0.05, ^#^*p*<0.05 by Bonferroni’s posttest between groups indicated on the bar graph.

To determine whether histone H1.2 is required for PAR translocation from the nucleus to the cytosol, histone H1.2 was knocked down with CRISPR/Cas9 and 2 different sgRNAs. Histone H1.2 sgRNA#1 and sgRNA#2 knockdown expression of histone H1.2 by approximately 50% (Fig. S2A). Both histone H1.2 sgRNA#1 and sgRNA#2 significantly reduce the translocation of PAR from the nucleus to the cytosol in response to NMDA or OGD as monitored by immunohistochemistry (Fig. 2A-D). These findings were confirmed by subcellular fractionation and immunoblot analysis (Fig. S2B-E). Accompanying the reduction in PAR translocation is a significant reduction in cell death induced by NMDA or OGD (Fig. 2E-G). The shRNA knockdown of histone H1.2 with two different shRNAs reduced histone H1.2 expression by greater than 50% (Fig. S3A). Consistent with the CRISPR/Cas9 knockdown of histone H1.2, shRNA knockdown of histone H1.2 prevents PAR translocation following NMDA or OGD treatment (Fig. S3B-I). Additionally, shRNA knockdown of histone H1.2 significantly reduces NMDA- and OGD-induced cell death (Fig. S3J-K). To confirm that histone H1.2 is required for nuclear PAR translocation to the cytosol, isolated nuclei from histone H1.2 knockout, heterozygote and wild type cortical tissue were exposed to N-methyl-N’-nitro-N-nitrosoguanidine (MNNG), a DNA alkylating agent that potently activates PARP-1. MNNG induces PAR translocation from wild type nuclei but not histone H1.2 knockout or heterozygote nuclei (Fig. S4A-B).

**Figure 2.**
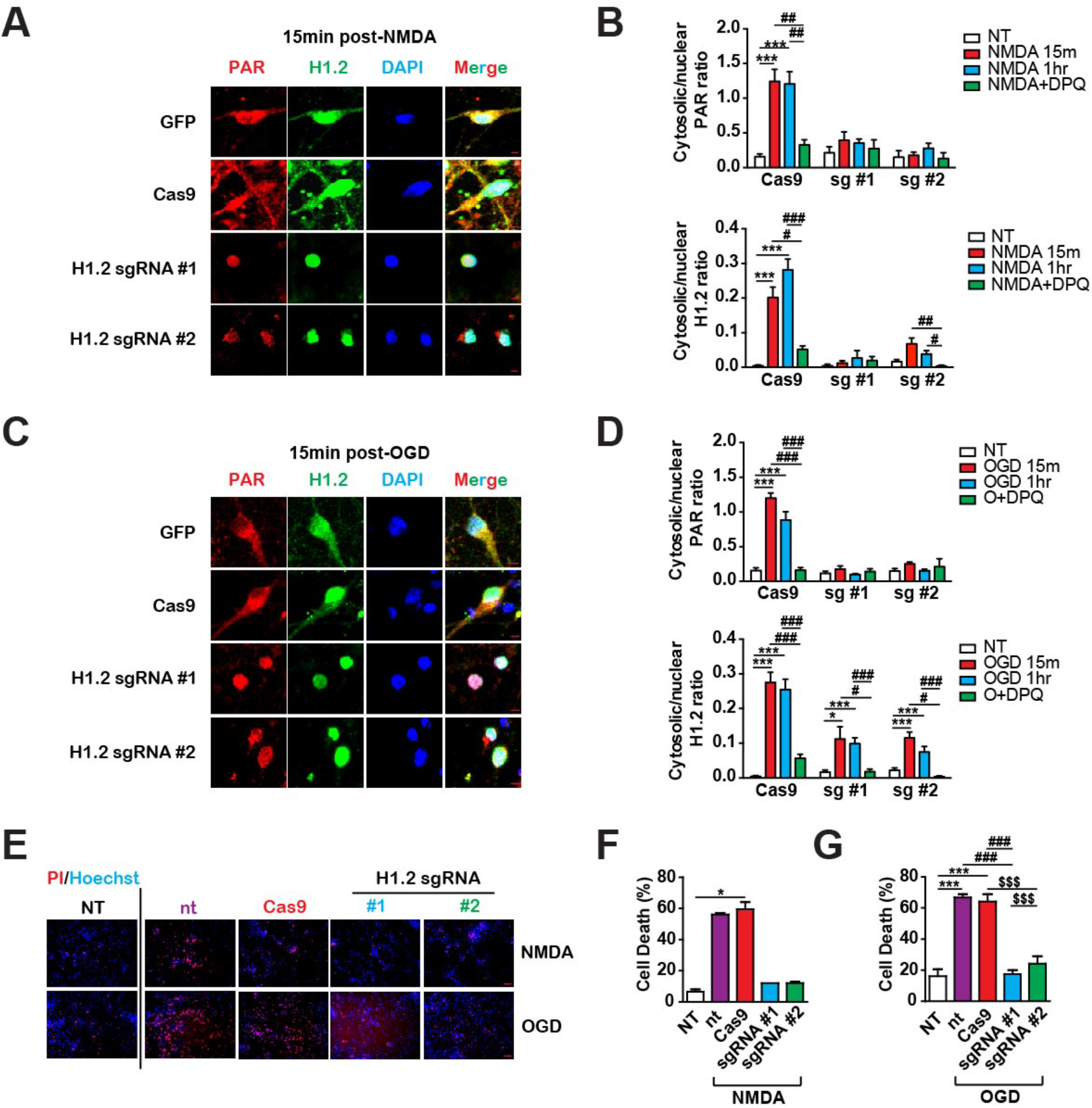
H1.2 deletion reduces PAR/H1.2 translocation and neuronal death induced by NMDA and OGD. (**A-D**) Representative photomicrographs showing human cortical neurons transduced with lentiviruses carrying GFP, or Cas9, or Cas9 and sgRNAs #1 and #2 targeting H1.2. Cells were fixed with 4% PFA 15 minutes after NMDA (**A**) or OGD (**C**) challenge, stained with anti-PAR antibody (red), anti-H1.2 antibody (green) and DAPI (blue). The merged panel is a composite of the red, green and blue channels. The cytosolic and nuclear levels of H1.2 and PAR are quantified, and the ratios (translocation) are expressed as mean ± s.e.m of 15 independent cells from 2-3 independent experiments. *P*<0.001 by one-way ANOVA for gene, treatment, and interaction of both PAR and H1.2 translocations in NMDA (**B**) and OGD (**D**) experiments, \**p*<0.05, \*\*\**p*<0.001, ^#^*p*<0.05, ^##^*p*<0.01, ^###^*p*<0.001 when compared between indicated groups by Bonferroni’s posttest. sg #1, sgRNA #1; sg #2, sgRNA #2. Scale bar = 10 µm. (**E-G**) Human cortical neurons are transduced with lentiviruses carrying Cas9 only or Cas9 and two sgRNAs targeting H1.2, #1 and #2 respectively. Percentages of cell death were assessed by PI/Hoechst stain 24 hrs after 30 minutes of 500 μM NMDA or 2 hrs of OGD (**E**) insults, quantified and expressed as mean ± s.e.m of 2 or 4 independent experiments. *P*<0.001 by one-way ANOVA for both NMDA (**F**) and OGD (**G**) experiments, ^###^*p*<0.001, \**p*<0.01, \*\*\**p*<0.001, ^$$$^*p*<0.001 when compared between indicated groups by Bonferroni’s posttest. NT, no transduction and no treatment; nt, no transduction; Cas9, Cas9 only. Scale bar = 100 µm.

To ascertain whether the export of histone H1.2 from the nucleus to the cytosol is required for PAR translocation and cell death, the antibiotic nuclear export inhibitor leptomycin B ^41^ was utilized. Leptomycin B treatment prevents both PAR and histone H1.2 translocation from the nucleus to the cytosol induced by NMDA or OGD (Fig. 3A-D). In a similar manner, Leptomycin B prevents both NMDA- and OGD-induced cell death (Fig. 3E-F).

**Figure 3.**
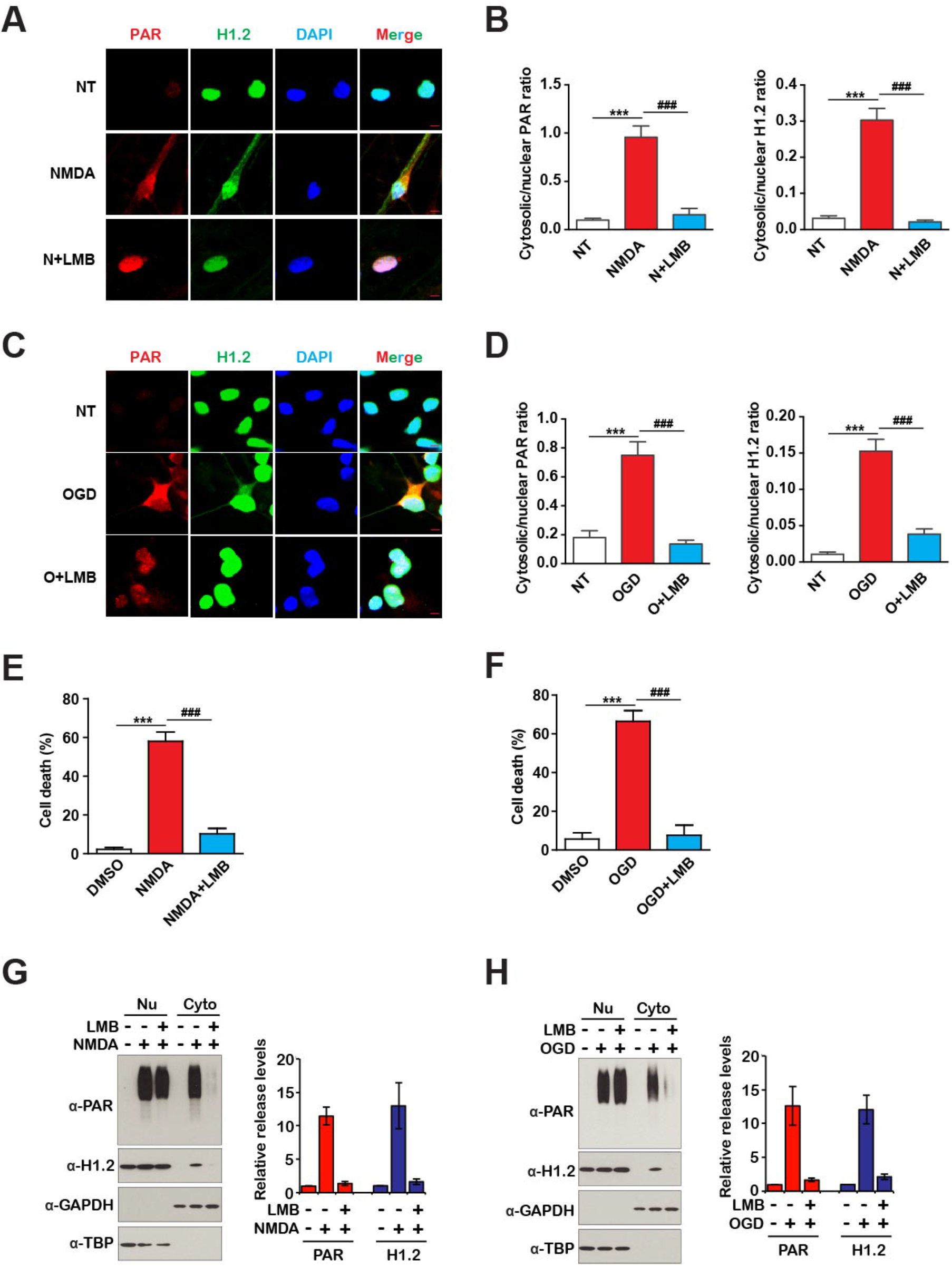
Leptomycin B (LMB) reduces PAR/H1.2 translocation and neuronal death induced by NMDA and OGD. (**A-D**) Representative photomicrographs showing human cortical neurons without any treatment (NT), with or without treatment of Leptomycin B (LMB) and NMDA (**A**) or OGD (**C**) insults. Cells were fixed with 4% PFA 15 minutes after 30 minutes 500 μM NMDA (**A, B**) or 2 hrs OGD (**C, D**) challenge, stained with anti-PAR antibody (red), anti-H1.2 antibody (green) and DAPI (blue). The merged panel is a composite of the red, green and blue channels. The cytosolic and nuclear levels of H1.2 and PAR are quantified and the ratios (translocation) are expressed as mean ± s.e.m of 15 independent cells from 2-3 independent experiments. *P*<0.001 by one-way ANOVA of both PAR and H1.2 translocations in NMDA (**B**) and OGD (**D**) experiments, \*\*\**p*<0.001, ^###^*p*<0.001 when compared between indicated groups by Bonferroni’s posttest. N+LMB, NMDA + Leptomycin; O+LMB, OGD + Leptomycin. Scale bar = 10 µm. (**E, F**) Human cortical neurons are treated with DMSO, and NMDA (**E**) or OGD (**F**) insults with or without LMB. Percentages of cell death were assessed by PI/Hoechst stain 24 hrs after, quantified and expressed as mean ± s.e.m of 3 independent experiments. *P*<0.001 by one-way ANOVA for NMDA and OGD experiments respectively,\*\*\**p*<0.001, ^###^*p*<0.001 when compared between indicated groups by Bonferroni’s posttest. (**G-H**) Representative immunoblots showing cytosol (**Cyto**) and nuclear (**Nu**) fraction of human cortical neurons 1 hr after challenged by NMDA (**G**) or OGD (**H**) and with or without leptomycin B. Blots were probed with anti-PAR, anti-H1.2, anti-TBP and anti-GAPDH antibodies. The releases of PAR and H1.2 from human cortical nuclei after NMDA or OGD are quantified and the PAR/GAPDH ratios (PAR release) or H1.2/GAPDH ratios (H1.2 release) shown are from 3 independent experiments, \**p*<0.05, \*\**p*<0.005 by one-way ANOVA with Bonferroni’s posttest. Nu, nucleus; Cyto, cytoplasm.

Iduna (RNF146) is a PAR dependent E3 ubiquitin ligase that targets PARylated or PAR binding proteins for ubiquitin proteosomal-dependent degradation ^42, 43^. Previous studies indicate that histones, including histone H1 bind to Iduna in a PAR dependent manner ^43^. To determine whether Iduna ubiquitinates PARylated histone H1.2, an *in vitro* ubiquitination assay was performed. Iduna ubiquitinates PARylated histone H1 and histone H1.2 while Iduna YRAA fails to ubiquitinate histone H1.2 (Fig. S5A). The effect of Iduna on histone H1.2 mediated nuclear to cytosolic translocation of PAR was evaluated. Overexpression of wild type Iduna prevents the translocation of histone H1.2 and PAR following NMDA or OGD exposure similar to inhibition of PAR formation by DPQ, while the catalytically null variant Iduna C60A, or the PAR binding mutant Iduna YRAA are ineffective (Fig. 4A-E, G-K). In addition, overexpression of Iduna, but not Iduna C60A or YRAA mutants, significantly reduces NMDA and OGD induced human cortical neuron cell death (Fig. 4F, L). Consistent with the observation that Iduna ubiquitinates PARylated histone, Iduna overexpression reduces the expression levels of cytosolic histone H1.2 and PAR following NMDA or OGD while the Iduna C60A or YRAA mutants have minimal effects (Fig. S5B-E).

**Figure 4.**
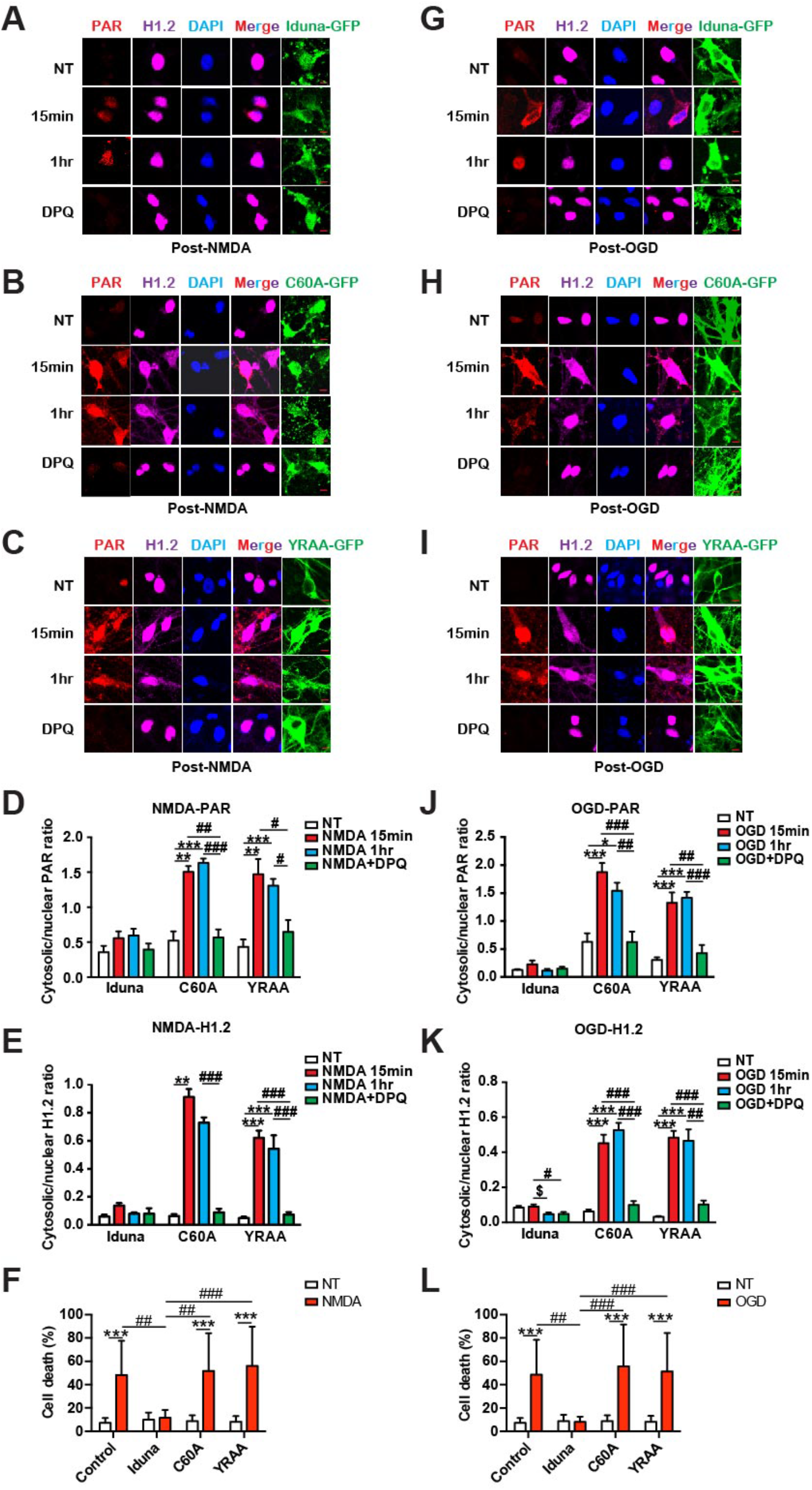
Overexpression of Iduna but not its mutants reduces PAR and H1.2 translocation induced by NMDA and OGD. (**A-F**) Representative photomicrographs showing human cortical neurons are transduced with lentiviruses carrying Iduna-GFP (**A**) or C60A-GFP (**B**) or YRAA-GFP (**C**) respectively. Cells were fixed with 4% PFA 15 minutes and 1hr after NMDA challenge with or without DPQ treatment, stained with anti-PAR antibody (red), anti-H1.2 antibody (magenta) and DAPI (blue). The merged panel is the composite of red, magenta and blue channels. NT, no treatment. The cytosolic and nuclear levels of PAR (**D**) and H1.2 (**E**) in (**A**) to (**C**) are quantified and the ratios (translocation) are expressed as mean ± s.e.m of 15 independent cells from 3 independent experiments. *P*<0.001 by one-way ANOVA for gene, treatment, and interaction of both PAR and H1.2 translocations, \*\**p*<0.01, \*\*\**p*<0.001, ^#^*p*<0.05, ^##^*p*<0.01, ^###^*p*<0.001 when compared between indicated groups by Bonferroni’s posttest. (**F**) Human cortical neurons transduced with lentiviruses carrying Iduna-GFP or C60A-GFP or YRAA-GFP were challenged with NMDA and assessed for death. Quantitative data is shown in (**F**) as mean ± s.e.m. of PI/Hoechst positive cell ratio by staining of three independent cultures and experiments. *P*<0.05, or *P*<0.001 by one-way ANOVA for gene, treatment, and interaction, *** *p*<0.001, ^##^*p*<0.01, ^###^*p*<0.001, when compared between indicated groups by Bonferroni’s posttest. Scale bar = 10 µm. (**G-L**) Same as above, but with OGD challenges. ^#^ or * or ^$^ *p*<0.05, ^##^ *p*<0.01, ^###^ or \*\*\**p*<0.001 when compared between indicated groups by Bonferroni’s posttest. Scale bar = 10 µm.

The C-terminal domain of histone H1 family proteins is critical for their protein binding and subcellular localization ^44^. In addition, the C-terminal domain of histone H1.2 is responsible for its actions in mediating cell death ^45, 46^. Two different C-terminal mutants of histone H1.2 were examined. C1 lacks amino acids 203-213 and C2 lacks amino acids 191-213 (Fig. 5A). Both the C1 and C2 mutants were expressed in human cortical cultures in which histone H1.2 was knocked down (Fig. S6A). In cultures expressing either the C1 or C2 mutant, histone H1.2 and PAR fail to translocate out of the nucleus (Fig. 5B-E). Since lysine 213 is known to be ribosylated on histone H1.2 (ADD REF 218926?), a lysine to alanine mutant (K213A) was examined in human cortical cultures in which histone H1.2 was knocked down (Fig. S6A). Histone H1.2 K213A has no effect on histone H1.2 and PAR translocation following NMDA or OGD stimulation (Fig. 5B-E). The C1 and C2 histone H1.2 mutants reduce NMDA- or OGD-induced cell death in histone H1.2 knockdown cultures while wild type histone H1.2 or histone H1.2 K213A restore neurotoxicity (Fig. 5F-H). A PAR binding assay indicates that histone H1.2 K213A and the C2 mutant bind PAR while the C1 histone H1.2 mutant is devoid of PAR binding (Fig. S6B). Histone H1.2 K213A has comparable ribosylation as compared to wild type histone H1.2 while the C1 and C2 mutants have reduced ribosylation (Fig. S6C). These results taken together suggest that both PAR binding and ribosylation of histone H1.2 are involved in the translocation of PAR from the nucleus to the cytosol.

**Figure 5.**
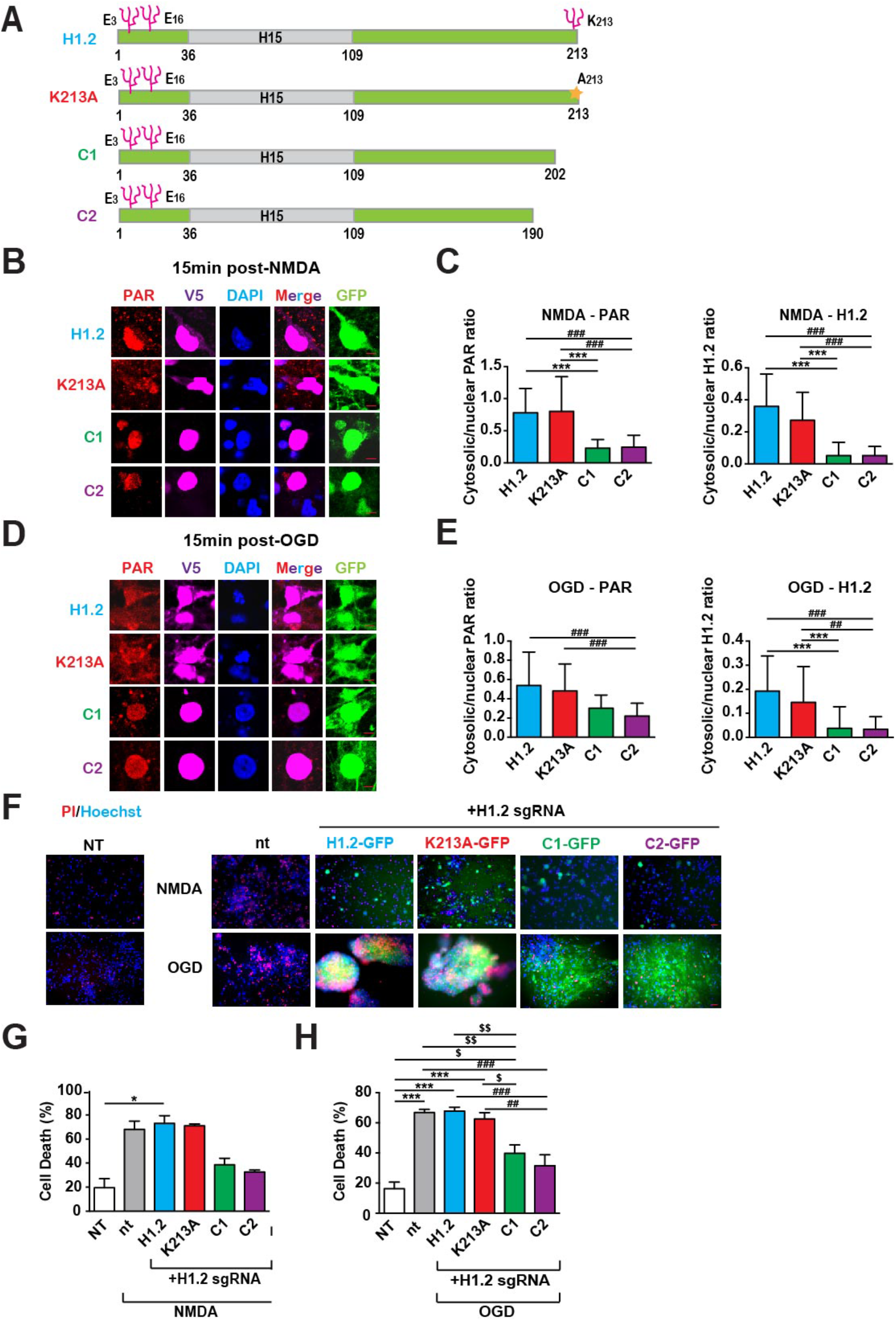
H1.2 carboxyl terminus is involved in Parthanatos and PAR/H1.2 translocation upon NMDA and OGD stimuli. (**A**) Schematic showing previously identified parylation site K213 ^52^, mutant K213A and two carboxyl tail truncated forms (C1, C2) of human histone H1.2 protein. (**B-E**) Representative photomicrographs showing human cortical neurons transduced with lentiviruses carrying Cas9, GFP and sgRNAs targeting H1.2, and overexpressed one of the following V5-tagged proteins: H1.2 (wild type), K213A, C1, or C2. Cells were fixed with 4% PFA 15 minutes after NMDA (**B**) or OGD (**D**) challenge, stained with anti-PAR antibody (red secondary antibody), anti-V5 antibody (magenta secondary antibody) and DAPI (blue). The merged panel is a composite of the red, magenta and blue channels. The cytosolic and nuclear levels of H1.2 and PAR are quantified and the ratios (translocation) are expressed as mean ± s.e.m of 20 cells from 3 independent experiments. *P*<0.001 by one-way ANOVA for both PAR and H1.2 translocations in NMDA (**C**) and OGD (**E**) experiments, ^##^*p*<0.01, ^###^ or \*\*\**p*<0.001, when compared between indicated groups by Bonferroni’s posttest. Human cortical neurons are transduced with lentiviruses carrying Cas9 and sgRNAs targeting H1.2, and overexpress one of the following GFP-tagged proteins: H1.2 (wild type), K213A, C1, or C2. After NMDA or OGD insult, PI and Hoechst are used to assess human neuronal death. Scale bar = 10 µm. (**F-H**) Percentages of cell death were assessed by PI/Hoechst stain 24 hrs after NMDA or OGD insults (**F**), quantified and expressed as mean ± s.e.m of 4 independent experiments. *P*<0.001 by one-way ANOVA for both NMDA (**G**) and OGD (**H**) experiments, \**p*<0.05, \*\*\**p*<0.001, ^##^*p*<0.01, ^###^*p*<0.001, and ^$^*p*<0.05, ^$$^*p*<0.01, when compared between indicated groups by Bonferroni’s posttest. NT, no transduction and no treatment; nt, no transduction. Scale bar = 100 µm.

To query the relevance of these findings in an *in vivo* setting, the middle cerebral artery occlusion (MCAO) reperfusion injury model was chosen as it has a strong parthanatos component ^5, 47^. Prior to MCAO, cortical cultures from histone H1.2 heterozygote, knockout mice and wild type mice were subjected to NMDA or OGD. Histone H1.2 heterozygote and knockout cultures are resistant to NMDA or OGD neurotoxicity (Fig. S7). Next, male histone H1.2 heterozygote, knockout and age matched littermate controls were subjected to 60 minutes of MCAO followed by reperfusion. Infarct volume was assessed 24 hrs and 7 days later. Infarct volume is substantially reduced in both histone H1.2 heterozygote and knockout mice compared to littermate controls 24 hrs post MCAO (Fig. 6A-G) and 7 days post MCAO (Figure 6L-N). Histone H1.2 heterozygote and knockout mice show better recovery in motor deficits than wildtype littermates at day 3 and/or day 7 post MCAO (Figure 6O-Q). Taken together these data indicate an important role for histone H1.2 in neuronal injury following stroke.

**Figure 6.**
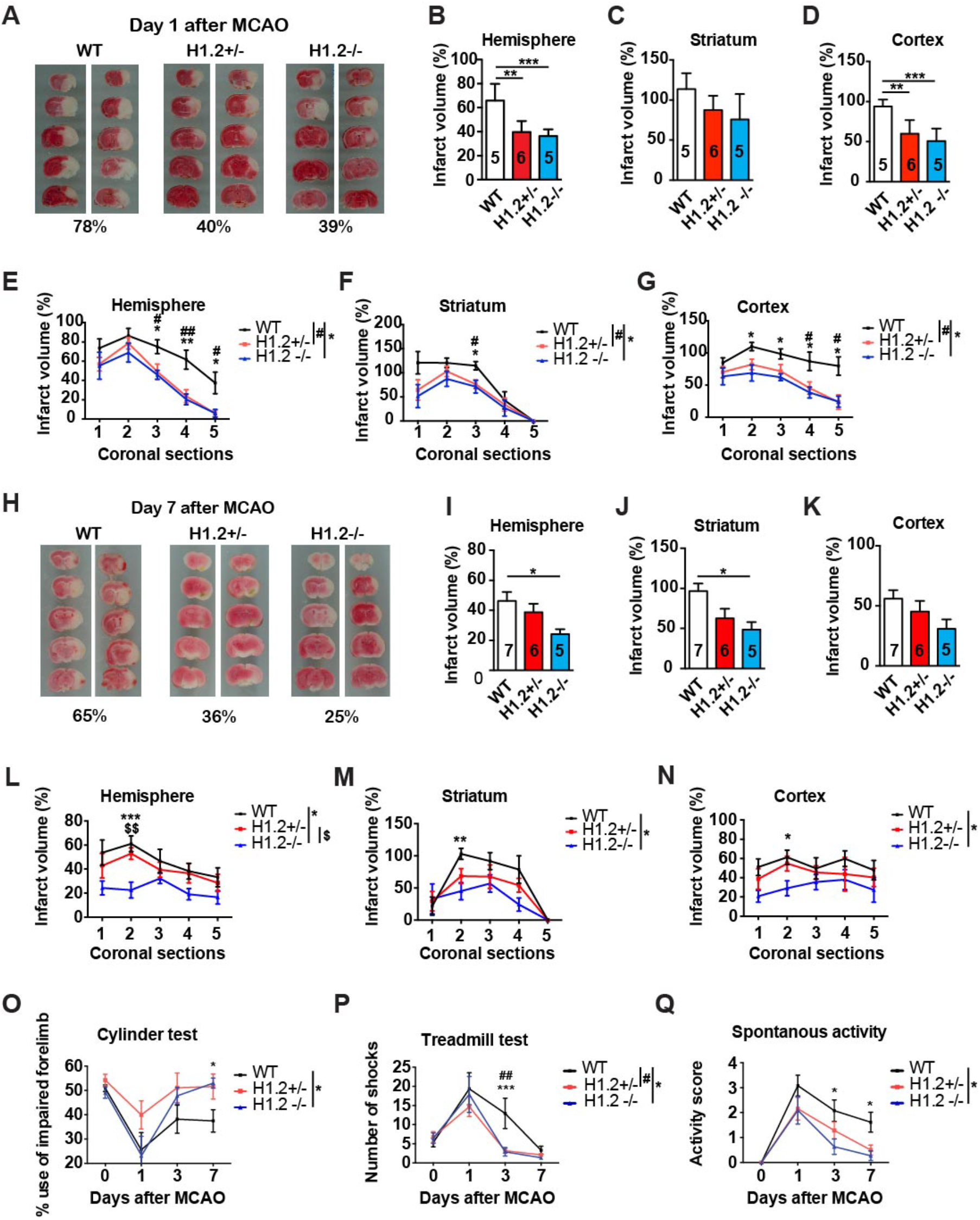
H1.2 partial or complete deletion reduces mice cortical neuronal death induced by NMDA or OGD, and deficits in mouse MCAO model. (**A-N**) Examples of TTC stained wild type and H1.2 knockout mice brain slices (both sides) obtained 24 hrs (**A**) or 7 days (**H**) after 1 hr transient MCAO. The pooled data of total (**B, C, D; I, J, K**) or coronal section (**E, F, G; L, M, N**) brain infarct volume percentages are shown as mean ± s.e.m. of 5 or 6 animals as indicated for whole hemisphere (**B, I**), striatum (**C, J**) and cortex (**D, K**). *P*<0.001 by one-way ANOVA, \*\*\**p*<0.001, \*\**p*<0.01, when compared to wild type (WT) animals by Bonferroni’s posttest. *P*<0.001 by one-way ANOVA across genotypes and sections of infarct volume data by sections (**E-G, L-N**), ^#^ or \**p*<0.05, ^##^ or \*\**p*<0.01, when compared to H1.2+/-, and ^#^ or * *p*<0.05, ^$^*p*<0.05, ^$$^*p*<0.01 when compared to H1.2-/- animals by Bonferroni’s posttest. **(O-Q)** The pooled data of cylinder test (**O**), treadmill test (**P**) and spontaneous activity (**Q**) are shown as mean ± s.e.m. of 7-9 animals of each genotype with day 7 endpoints. *P*<0.001 by one-way ANOVA (total infarct volume), \*\**p*<0.01, when compared to wild type (WT) animals by Bonferroni’s posttest. *P*<0.001 by one-way ANOVA for genotypes and sections of the bottom panels, \**p*<0.05, \*\*\**p*<0.001, when WT compared to H1.2+/-, and ^#^*p*<0.05, ^##^*p*<0.01 when WT compared to H1.2-/- animals by Bonferroni’s posttest.

## Discussion

The major finding of this study is that histone H1.2 is required for nuclear PAR translocation to the cytosol and parthanatos both *in vitro* and *in vivo*. PARP-1 activation leads to PAR formation in response to a toxic dose of NMDA or prolonged OGD ^11, 48, 49^. Both PAR and histone H1.2 translocation from the nucleus to the cytosol occurs within 15 minutes of either NMDA or OGD exposure. Inhibition of PARP-1 by DPQ, blockade of nuclear export by Leptomycin B or expression of Iduna is sufficient to reduce PAR and histone H1.2 translocation and neuronal cell death. The C-terminus of histone H1.2 is critical for the translocation of histone H1.2 and PAR in parthanatos. Knockdown of histone H1.2 protects against NMDA or OGD neuronal cell death in human cortical cultures. Knockout of histone H1.2 in mice reduces injury following NMDA or OGD induced cell death in murine cortical cultures or MCAO *in vivo*.

Of the histone 1 family members, histone H1.2 plays an active role in different forms of cell death ^38, 39, 50^. The C1 and C2 regions of histone H1.2 are divergent from other histone H1 family members ^35^. This divergence may account for histone H1.2 cell death promoting properties. Histones are PARylated, as well as, PAR binding proteins. Our mutational analysis suggests that both PARylation and PAR binding of histone H1.2 are required for PAR translocation and parthanatos. Future studies will be required to determine the relative contributions of ribosylation and PAR binding in these processes.

These results are consistent with the notion that PAR is a death signal when in the cytosol in parthanatos ^11^. Over expression of the PAR degrading enzyme, PARG, protects against parthanatos ^16^, while knockdown or knockout of PARG exacerbates PARP-1 dependent cell death ^51^. The protection afforded by ARH3 suggests that free PAR polymer but not ribosylated proteins are the death signal ^20^. Thus, PAR binding to histone H1.2 may be more important than ribosylation.

In summary, histone H1.2 is a major contributor to parthanatos by carrying PAR out of the nucleus to the cytosol initiating the transition from the nucleus to the mitochondria in in parthanatic death cascade. Strategies aimed at interfering with PAR signaling may offer novel therapeutic opportunities to treat disorders characterized by overactivation of PARP-1.

## Acknowledgements

The authors thank the Dawson laboratory and Jin-Chong Xu and Shaida Andrabi for helpful discussion and Josh Ostovitz, Rong Chen, Li Chen and Juehua Zhu for technical assistance. This work was supported by grants from the National Institutes of Health (NIH) NIH/ NIDA DA000266 and NIH/NINDS R01 NS067525, R37 NS067525 and NS38377 to T.M.D. and V.L.D. T.M.D. is the Leonard and Madlyn Abramson Professor in Neurodegenerative Diseases. The authors acknowledge the joint participation by the Adrienne Helis Malvin Medical Research Foundation through its direct engagement in the continuous active conduct of medical research in conjunction with The Johns Hopkins Hospital and the Johns Hopkins University School of Medicine and the Foundation’s Parkinson’s Disease Program M-2016.

## Author Contributions

Conceptualization, J.F., T.M.D., V.L.D.; Methodology, J.F., B.G.K., T.-I.K., J.-C.X.; Validation, J.F., B.G.K. T.-I.K.; Formal Analysis, J.F., B.G.K., T.-I.K., A.B., Investigation, J.F., B.G.K., T.-I.K., J.-C.X., J.R.O., J.R., A.B., H.K.,; Resources, V.L.D., T.M.D.; Writing-Original Draft, J.F, T.M.D., V.L.D.; Writing, Reviewing & Editing, J.F, T.M.D., V.L.D.; Visualization, J.F., B.G.K., T.-I.K., J.-C.X.; Supervision, T.M.D., V.L.D.; Project Administration, T.M.D., V.L.D.; Funding Acquisition, T.M.D., V.L.D.

## Methods

### hESCs culture and neural differentiation

hESC lines H1 were maintained and differentiated into cortical neurons following published lab protocols ^40^. The human H1 cortical neuron cultures are used at 2 months after induction of differentiation when neurons are relatively mature and show responses to toxic NMDA or OGD stimulation ^40^. All experiments using H1 hESCs cells are conducted in compliance with the policy of the JHU SOM and are oversighted by the JHU Institutional Stem Cell Research Oversight (ISCRO) Committee.

### Primary neuronal culture preparation

Primary cortical cell cultures of wild type or H1.2 knockout C57BL/6 littermate mice were prepared from gestational day 14-15 embryos, and maintained in neurobasal medium with B27 supplements until day 12-14 *in vitro*. On the day of experiments, neurons represent 90% of the cells in the culture.

### NMDA treatment

As described in ^50, 53^, either mouse or human neurons were washed with control salt solution (CSS) containing 120 mM NaCl, 5.4 mM KCl, 1.8 mM CaCl_2_, 25 mM Tris-HCl and 15 mM D-glucose (pH 7.4) before treatment. To induce toxicity, 500 µM NMDA and 10 µM glycine in CSS solution was applied to the mouse neuron cultures for 15 minutes or to the human neuron cultures for 30 minutes. Then the cultures were washed once with CSS and replaced with saved conditional medium. No treatment controls were incubated with the CSS alone for the same length of time.

### Oxygen Glucose Deprivation (OGD)

As described in ^53^, glucose-free HBSS was bubbled with OGD gas (5% CO_2_, 10% Hydrogen, and 85% N_2_, Airgas Ltd. USA) for 30 min to first remove dissolved oxygen. Neuronal cultures were then washed once with this pre-bubbled glucose-free media and then immediately incubated in a hypoxia incubator with oxygen sensor and monitor (Biospherix Ltd. USA) in the same media for 1 hr (mouse neuron cultures) or 2 hrs (human neuron cultures). Neuron cultures were then re-fed with saved normal conditional media and maintained under normal culture condition. The no treatment controls were kept in normal growth media and normal incubators.

### Nuclei isolation and MNNG treatment

Nuclei of cortical tissues from wild type (WT), histone H1.2 knockout heterozygous (Het) and homozygous (KO) mice are isolated using the Nuclei Isolation ‘Kit Nuclei EZ Prep’ (NUC101, Sigma). Then the nuclei are treated with 500 µM MNNG with fresh-made NAD+ (N0632, Sigma) in PBS for 15 minutes. Then the nuclei are collected and resuspended in PBS. Fifteen minutes after treatment, the nuclei buffer (PBS) and nuclei lysates are collected separately and forwarded to western blot analysis.

### Antibodies

The following antibodies were used at dilutions suggested by the manufacturer’s instructions. Primary antibodies include: rabbit anti-H1.2 (ab17677, Abcam, RRID:AB_2117984), mouse anti-eGFP (MA1-952, Invitrogen, RRID: AB_889471), mouse anti-V5 (R960-25, Invitrogen, RRID: AB_2556564), mouse anti-FLAG (M2, F1804, Sigma-Aldrich), mouse and rabbit anti-poly(ADP-ribose) antibodies (4335-MC-100, RRID:AB_2572318 and 4336-BPC-100, Trevigen,), human anti-poly(ADP-ribose) antibodies (Dawson laboratory, Human #19), anti-poly(ADP-ribose) polymerase (556494, RRID:AB_396433, mouse, BD bioscience; 9542, RRID:AB_2160739, rabbit, Cell signaling Technology), rabbit anti-TBP (#8515, Cell Signaling,), rabbit anti-alpha-Tubulin (#2144, Cell signaling), horse radish peroxidase (HRP) conjugated mouse anti-β-actin (#5125, Cell Signaling), HRP conjugated mouse anti-GAPDH (#3683, Cell Signaling,), rabbit anti-Iduna (ab201212, Abcam). Secondary antibodies include: HRP conjugated goat anti-mouse IgG (#7076, Cell signaling), HRP conjugated goat anti-rabbit IgG (#7074, Cell signaling), Alexa Fluor 488 conjugated goat anti-mouse or rabbit IgG (H+L) (Invitrogen), Alexa Fluor 594 conjugated goat anti-mouse or rabbit IgG (Invitrogen), Alexa Fluor 647 conjugated goat anti-mouse or rabbit IgG (Invitrogen).

### Immunoprecipitation

Immunoprecipitation (IP) used antibodies to H1.2, V5, PAR (Trevigen and human #19) or magnetic beads conjugated with anti-Flag antibody (Sigma). Protein G–agarose beads (GE Healthcare) or magnetic beads were washed twice before immunoprecipitation.

Human neuronal cells were lysed in RIPA buffer (50 mM Tris-HCl, pH 7.4, 150 mM NaCl, 1 mM EDTA, 1 mM EGTA, 1% NP-40, supplemented with a complete protease inhibitor cocktail) and passed through 26G syringe needle for 5 times. After 30 minutes incubation on ice, samples are centrifuged at 15,000 X g for 15 minutes at 4°C. The supernatants were used for immunoprecipitation by incubation for 2 hrs or overnight at 4°C with antibodies or IgG of the same species and beads. After 5 times washing with RIPA buffer, the proteins on the beads were eluted with 4X protein loading buffer (Bio-Rad) and heated at 97° C for 5 minutes. Then the IP samples and saved cell lysates (input) were subjected to western blot analysis. Protein levels were quantified using a BCA protein assay kit (Pierce). Brain tissue lysates (40 μg) or IP’ed samples were electrophoresed on SDS-PAGE gels and transferred to nitrocellulose membranes. Membranes were blocked with 5% skim milk in TBS-T and incubated with primary antibodies. After HRP-conjugated secondary antibodies incubation, the immunoblot signal was detected using chemiluminescent substrates (Thermo Scientific).

### Subcellular fractionation

Ventral midbrain tissues from mice or ice cold PBS-washed SH-SY5Y cells were homogenized in a hypotonic buffer (10 mM HEPES, pH 7.5, 10 mM KCl, 1.5 mM MgCl2, 1 mM EDTA, 1 mM EGTA, 1 mM DTT, and 0.1% NP-40) supplemented with a complete protease inhibitor cocktail and passed through 26G syringe needle for 3 times. After incubation on ice for 20 minutes with 10 seconds vortexing two times, the lysates were centrifuged at 800 *g* for 5 minutes. The supernatant was used as a post-nuclear fraction and the pellet was washed with the same buffer twice and used as a nuclear fraction for subsequent western blot analysis (Loading for western blot: 2% of PN fraction and 8% of Nu fraction).

### Immunocytochemistry, immunohistochemistry, imaging and quantification

Cultured cells were washed in PBS and fixed in 4% paraformaldehyde for 15 min. Attached RONAs and human cortical assemblies were sectioned (25 μm) by cryostat (CM3050; Leica, Nussloch, Germany) and collected on SuperFrost Plus glass slides (Roth, Karlsruhe, Germany). Cells and sections were blocked with blocking buffer containing 10% v/v donkey serum and 0.2% v/v Triton X-100 in PBS. Primary antibodies diluted in blocking buffer were incubated overnight at 4°C, washing three times in blocking buffer, treatment with secondary antibody (Invitrogen) were applied for 1 hr and washing three times in blocking buffer. After staining, coverslips were mounted on glass slides and sections were coverslipped using prolong gold antifade reagent (Invitrogen).

### Cell Death Assessment

Two to three month old human cortical neurons were treated with NMDA or OGD at various indicated time duration and doses. Percent of cell death was determined by the staining with 5 µM Hoechst 33342 and 2 µM propidium iodide (PI) (Invitrogen, Carlsbas, CA). Images were taken and counted by Zeiss microscope equipped with automated computer-assisted software (Axiovision 4.6, Zeiss). 20 µM Z-VAD (Sigma, V116), 20 µM NEC-1 (Sigma, N9037), 500 µM 3-MA (Calbiochem, 189490), 20 µM NPLA (Tocris, 1200), 30 µM DPQ (Enzo, ALX-270-21-M005), 500 nM AG-014699 (Selleckchem, S1098) 10 µM ABT888 (Active biochem, A-1003), 2 µM Olaparib (LC laboratories, O-9201) and 20 nM BMN673 (Selleckchem, S7048) were applied to evaluate the effect of different antagonists to known cell death signals. Following exposure of neuronal cultures to the various treatments, neuronal survival was quantified and presented as percent of cell death. Percent cell death was determined as the ratio of live-to-dead cells compared with the percent cell death in control wells to account for cell death attributable to mechanical stimulation of the cultures. Quantification of neuronal survival was determined by staining treated cultures with 5 µM Hoechst 33342 and 2 µM propidium iodide (PI) (Invitrogen, Carlsbad, CA) 1. Culture plates were placed on a mechanized stage of a Zeiss microscope and photomicrographs were collected by a blinded observer. The numbers of total and dead (PI positive) cells were counted by automated computer assisted software (Axiovision 4.6, Zeiss, Germany) 1. The raw counts are presented in an Excel file for generation of percent cell death and statistical analysis. Glial nuclei fluoresce at a lower intensity than neuronal nuclei and were gated out by the software program. At least two separate experiments using four separate wells were performed for all Nature Medicine doi:10.1038/nm.2387 data points.

### Constructs

Iduna complementary DNAs were cloned from mouse cDNA and sequenced. Iduna PCR products were cloned into the phCMV1-Xi vector (Gene Therapy Systems), pEGFP-C2 vector (Clontech), pCMV-Tag5 vector (Stratagene) and pGEX-6p vector (GE Health Care). Deletion mutants and YRAA mutants were constructed by PCR and were verified by sequencing.

### H1.2 knockout in human cortical neuronal culture using CRISPR/Cas9

PARP-1 sgRNAs (#1, GTGGCCCACCTTCCAGAAGC; #2, ATACCAAAGAAGGGAGT-AGC) were synthesized and subcloned into lenti-CRISPR vector (Addgene, pXPR_001, plasmid 49535). Lentiviral vectors were co-transfected into HEK293FT cells with the lentivirus packaging plasmids pVSVg and psPAX2 using FuGENE® HD. Supernatants containing virus were collected 48 and 72 hrs post transfection, passed through nitrocellulose filter (0.45 μm) and applied on cells in culture. At 8-12 weeks post-differentiation, human cortical neurons were transduced with Lenti-CRISPR lentivirus carrying control sgRNA or PARP-1 sgRNAs. Cells were then challenged with NMDA or OGD at 5 days after transduction, and cell death was assessed by PI/Hoechst stain 24 hrs later. Another batch of cells was collected at 5 days after transduction and forwarded to western blotting for the PARP-1 expression levels.

### Western blotting

SDS-PAGE and transfer were performed according to laboratory protocol with slight modification. In brief, cultured cells were lysed in RIPA buffer containing 1% Triton, 0.5% Na-deoxycholate, 0.1% SDS, 50 mM NaF, 10 mM Na_4_P_2_O_7_, 2 mM Na_3_VO_4_, and EDTA-free protease inhibitor mixture (Roche Diagnostics) in PBS (pH7.4). Protein extracts were separated by 4-12% SDS-PAGE. Proteins were transferred at a constant voltage of 80 V for 150 min at 4°C from the SDS to PVDF membranes. Membranes were then blocked with 5% non-fat milk and incubated with primary antibodies overnight at 4°C. Antibodies used were anti-PAR (Trevigen, 4336-APC-050), Actin (Cell Signaling, 5125), After washes with TBST (TBS with 0.1% Tween-20), membranes were incubated with HRP-conjugated secondary antibodies for 1 hr at RT. Immunoreactive bands were visualized by the enhanced chemiluminescent substrate (ECL, Pierce) on X-ray film and quantified using the image software TINA. Neuronal cultures were exposed to NMDA for 5 min. Cell lysates were subjected to centrifugation at 12,000 X g for 10 min at 4°C. The resulting supernatant was subjected to SDS-PAGE, and the separated proteins were transferred electrophoretically to a nitrocellulose membrane. The membrane was incubated with a Tris-buffered saline solution containing 5% nonfat milk and 0.1% Tween 20. The membrane was then incubated for 1 hr at room temperature with the indicated antibodies in a Tris-buffered saline solution containing 0.1% Tween 20 and subsequently with appropriate secondary antibodies conjugated with horseradish peroxidase (Amersham Biosciences). The immunoblots were visualized in X-ray films by an enhanced chemiluminescence method (Pierce, USA). Antibodies used include: anti-myc (Roche Applied Sciences USA), anti-PAR, anti-PARP-1 (BD Pharmigen USA), antiGFP, anti-COXIV (Abcam Inc Cambridge, MA), HRP conjugated anti-β-actin, anti-β-tubulin, and anti-biotin (Sigma, USA), anti-AIF (Epitomics, Burlingame, CA).

### Lentiviral preparations

We used Invitrogen ViraPower lentiviral packaging system and obtained high-titer viral preparations for effective transduction in primary neuronal cultures and for intra-striatal injections. For developing efficient shRNA lentiviruses, we subcloned a siRNA oligo directed to the coding region +556 – 576 of Iduna into a lentiviral Nature Medicine doi:10.1038/nm.2387 expression vector, cFUGw. The oligo was PCR amplified with primers flanked by PacI restriction sites. Following digestion and ligation, clones were selected and verified for the inserted sequence. The lentiviral construct co-expresses EGFP driven by the Ubiquitin C promoter, in addition to the mouse U6 Pol II promoter driving the shRNA. To control for off-target and nonspecific effects of shRNA, a shRNA against dsRed was used. The over expression lentiviral system was developed by removing the EGFP open reading frame from the cFUGw construct by a BamHI/XbaI digestion and replacing it with the cDNA of GFP-Iduna or GFP-Iduna YRAA. Near 100% neuron specific expression is observed, using either our over expression or RNAi lentiviral system. The cDNA of human Iduna was cloned from human MCF-7 cells mRNA by reverse transcription-PCR (RT-PCR) and then it was subcloned to pEGFP-C2 to create EGFP human-Iduna. The construction of cFUGW-EGFP human-Iduna, was performed by digesting the pEGFP-human-Iduna by BamHI/XbaI followed by subcloning into the same enzyme restriction sites of cFUGW. DNA sequences were verified by sequencing.

### Breeding and genotyping of H1.2 knock out mice

H1.2 knock out mice were genotyped by PCR of tail DNA with the following primers ^38^. The wild-type H1.2 allele (predicted size, 600 bp): PCF (5’-CTGCCACACCCAAAAAGG-3’) and PCR (5’-GAGCATAGAAGCCACCTACAAG-3’). The mutant H1.2 allele (predicted size, 400 bp): pGK2 (5’-GCTGCTAAAGCGCATGCTCCA-3’) and PCR (5’-GAGCATAGAAGCCACCTACAAG-3’). The Johns Hopkins Medical Institutions are fully accredited by the American Association for the Accreditation of Laboratory Animal Care. All research procedures were approved by the Johns Hopkins Medical Institutions Institutional Animal Care and Use Committee. All experiments were performed in accordance with the National Institutes of Health Guidelines and were approved by the institutional animal care and use committee.

### Middle cerebral artery occlusion

The middle cerebral artery was occluded in mice with a 7-0 monofilament (Ethicon) and confirmed by laser-Doppler flowmetry. After 60 min of occlusion, the filament was removed, and reperfusion was verified. At 24 h, the brain was collected and analyzed by an observer unaware of the genotype of the mouse. To occlude the middle cerebral artery, mice were anesthetized with 1.5–2% isoflurane and maintained at normothermic temperature. A 7-0 monofilament with an enlarged silicone tip was passed through the right internal carotid artery to the base of the middle cerebral artery. Occlusion was confirmed by laser-Doppler flowmetry with a probe placed on thinned skull over the lateral parietal cortex. After 60 min of occlusion, the filament was removed and reperfusion was verified. At 24 hr of reperfusion, the brain was harvested, sectioned into five coronal slabs, and stained with the vital dye, triphenyltetrazolium chloride. Infarct area was measured on the anterior and posterior surfaces of each slab and integrated to obtain infarct volume with correction for tissue swelling. The investigator performing the surgery and analyzing infarct size was unaware of the genotype of the mouse.

### Neurobehavioral activity in mice

Neurological deficits in spontaneous motor activity, treadmill, and cylinder test were evaluated at different times pre- and post MCAO by an observer blinded to the genotype of the mice. Spontaneous motor activity was evaluated for 5 min by placing the animals in a mouse cage for 5 minutes. A video camera was fitted on top of the cage to record the activity of a mouse in the cage. Neurological deficits of spontaneous motor activity were evaluated by an observer at a scale of 0-4 (0 no neurological deficit, 4 severe neurological deficit). The following criteria were used to score deficits: 0= mice appeared normal, explored the cage environment and moved around in the cage freely; 1 = mice hesitantly moved in the cage and did not approach all sides of the cage, 2 = mice showed postural and movement abnormalities and had difficulty approaching the walls of the cage, 3 = mice with postural abnormalities tried to move in the cage but did not approach the wall of the cage, 4 = mice were unable to move in the cage and stayed at the center. In the treadmill test, a mouse was placed at the middle of a treadmill (Panlab, Model LE8708) with electric bars at one end. The treadmill was turned on at setting 3 (Speed: 12cm/s, Shock intensity: 0.2mA) by the observer for 2 min. The number of shocks that the mouse received when it failed to keep up with the treadmill and dropped on the electric bar was recorded. The cylinder test was performed to assess the forelimb performance in mice. For this test, a transparent glass cylinder (9 cm in diameter and 15 cm in height) was placed in the center of a chamber containing two video cameras on opposite sides. A mouse was placed in the cylinder and the cameras on opposite sides were aligned at a straight axis with the cylinder to allow recordings of mouse forelimb movements on all sides of the cylinder. Recordings were evaluated by an observer blinded to the treatment and genotype of the animals. Forelimb use of the mouse was recorded for 10 minutes and analyzed according to the following criteria: (1) Ipsilateral (right) forelimb use (number of touches to the cylinder wall) independent of the left limb (2) Contralateral (left) forelimb use (number of touches to the cylinder wall) independent of the right limb (3) Simultaneous use of both limbs. The percent use of the contralateral (left) limb was quantified by subtracting contralateral fore paw touches from the total number of touches made by the mouse during the period of observation.

### Statistical analysis

Data are represented as mean ± s.e.m. Statistical analysis was performed using GraphPad Prism and noted in the text and figure legends. Information on sample size (n, given as a number) for each experimental group/condition, statistical methods and measures are available in all relevant figure legends. Unless otherwise noted, significance was assessed as *P*<0.05.

### Synthesis of [^32^P]-labeled poly(ADP-ribose) polymer (PAR)

3 ug of recombinant PARP1 was incubated in reaction buffer containing 100 mM Tris-Cl, pH 8.0, 10 mM MgCl2, 8 mM DTT, 10% glycerol, 23 ug calf thymus activated DNA, and 75uCi [^32^P]-labeled NAD for 30 min at 30°C. To collect automodified PARP1, 3 M CH3COONa and isopropanol were added to the sample, thereafter automodified PARP1 was precipitated by centrifugation at 10,000 g for 10 min. [^32^P] labeled PAR was alkali extracted from the automodified PARP-1 and purified through a dihydroxyboronyl BioRex (DHBB) column as described ^54^.

### *In vitro* ribosylation assay

1 ug of recombinant PARP1 was incubated in reaction buffer containing 100 mM Tris-Cl, pH 8.0, 10 mM MgCl2, 8 mM DTT, 10% glycerol and 25uCi [^32^P]-labeled NAD for indicated periods at 30°C. 23 ug of calf thymus activated DNA used if required. Reaction was stopped by addition of Laemmli buffer containing beta-mercaptoethanol and separated by SDS PAGE. Each gel was exposed to an imaging plate (Fuji film, BAS-MS) and the signal was visualized by a phosphorimager (GE, Typhoon FLA-9500).

### Electromobility gel shift assay

Each protein was incubated with purified [^32^P]-labeled PAR in binding buffer (50mM Tris pH7.5, 150 mM NaCl) at RT for 20 mins. Each sample was mixed with DNA loading dye (Thermo) and electrophoresed in 4-20% or 20% TBE PAGE (Invitrogen). Each gel was exposed to an imaging plate (Fuji film, BAS-MS) and signal was visualized by a phosphorimager (GE, Typhoon FLA-9500).

### *In vitro* ubiquitination assay

Recombinant E1, UbcH5c and His-ubiquitin were purchased from Boston Biochem. E1 (50 nM), UbcHs (50 nM) and each immunoprecipitated GFP-Iduna proteins were incubated with recombinant his-ubiquitin (200 mM) in reaction buffer containing 50 mM Tris-Cl, pH 8.0, 2.5 mM MgCl2, 2 mM DTT, 2 mM ATP at 37°C for 1 hr. Reaction was stopped by the addition of Laemmli buffer containing beta-mercaptoethanol and subjected to a following Western blot. Either His (Thermo) or GFP (Roche) or GST (Santa Cruz) antibody was used to detect ubiquitinated protein and free ubiquitin together.

## Supplemental Information

Extended data display items, video source files, key resource and statistical tables are available in the online version of the paper.

