## Supplemental Files for "Histone H1.2 Dependent Translocation of Poly (ADP-ribose) Initiates Parthanatos"

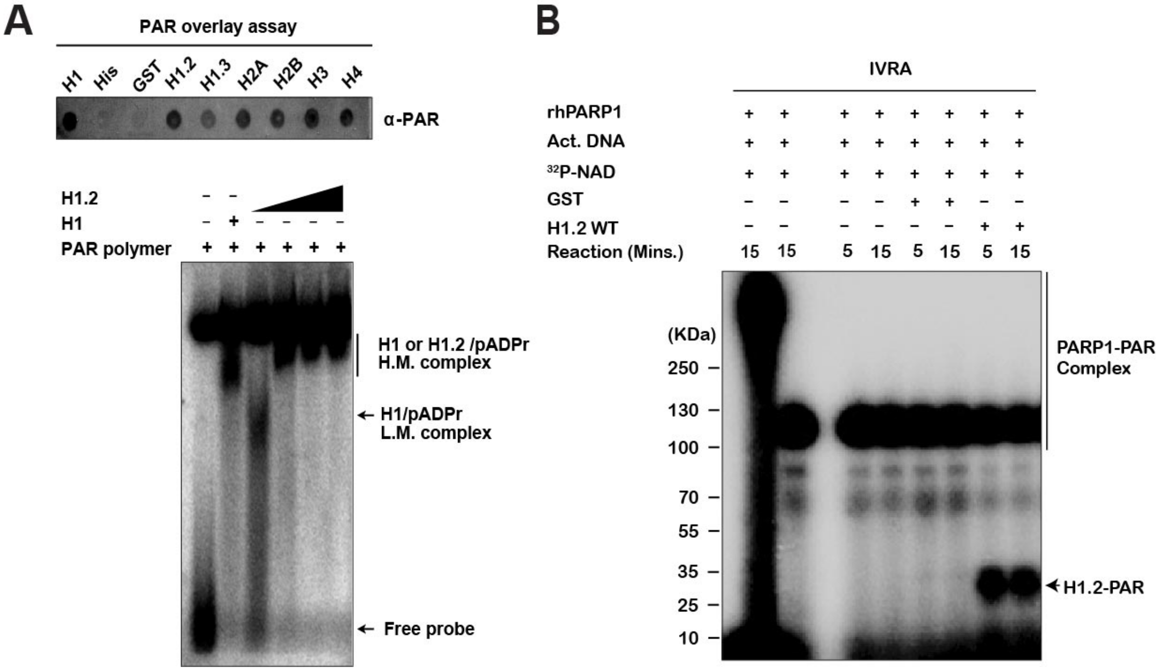


**Fig. S1. H1.2 binds PAR and can be ribosylated by PARP-1**

**(A**) PAR binding capacity of various histone protein were tested by PAR overlay assay. Histone H1 was used as a positive control. 6X His peptide and/or recombinant GST were used as a negative control.

**(B)** PAR binding capacity of H1.2 was tested by Electromobility shift assay. Recombinant H1 and an increasing amount of H1.2 were incubated with [$^{32}$P]-PAR and separated in 20% TBE PAGE. Both [$^{32}$P]-PAR bound signal and unbound free [$^{32}$P]-PAR signal were visualized by autoradiography.

**(C)** Ribosylation of H1.2 by PARP1 was tested by *in vitro* ribosylation assay. Hyperactivation of PARP1 in the presence of activated DNA showed inappropriate excessive signals followed by SDS PAGE (Lane1). The majority of auto-ribosylated PARP1 signal in the absence of activated DNA is detected in its corresponding molecular weight (Lane 2). Followed by *in vitro* ribosyation reaction, ribosylated-PARP1 (Lane4,

5) or -GST (Lane 6, 7) or -H1.2 (Lane 8, 9) were separated and visualized by autoradiography.

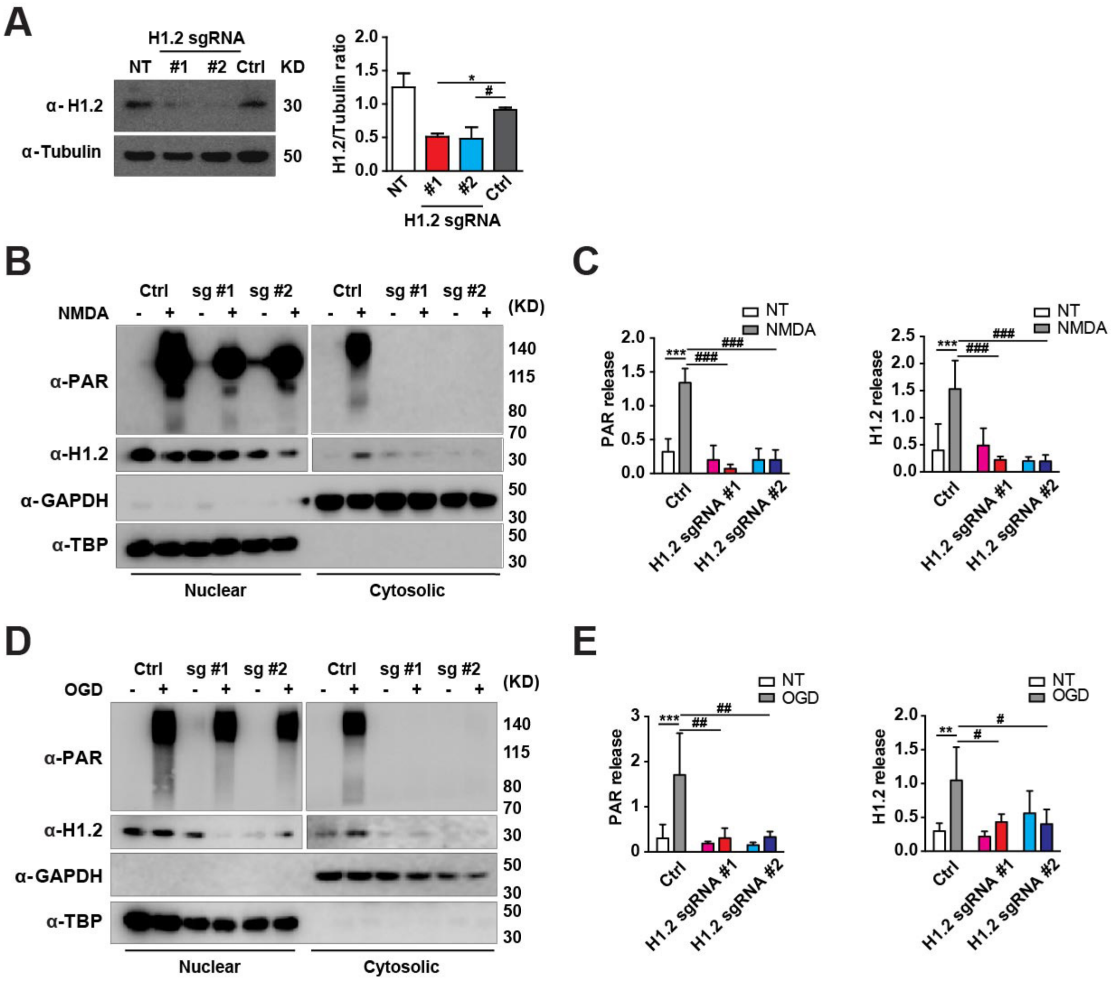


**Fig. S2. H1.2 sgRNA-mediated knock down reduces PAR/H1.2 translocation by NMDA and OGD.**

(**A**) Representative immunoblots showing the lysates of human cortical neurons transduced with lentiviruses carrying Cas9, and sgRNAs #1 and #2 targeting H1.2. Blots were probed with anti-H1.2 antibody and alpha-Tubulin. The protein levels of H1.2 are quantified and normalized to alpha-Tubulin protein levels. The ratios are expressed as mean ± s.e.m of 4 independent experiments. $P<0.01$ by one-way ANOVA, *$p<0.05$, when compared to the control (Ctrl, Cas9 only) group by Bonferroni's posttest. NT, no transduction.

(**B-E**) Representative immunoblots showing nuclear and cytosolic fractions of human cortical neurons transduced with lentiviruses carrying empty vector (Cas9), sgRNAs #1 (sg#1) and #2 (sg#2) targeting H1.2. Blots were probed with anti-PAR, anti-H1.2, anti-TBP and anti-GAPDH antibodies. The protein levels of PAR and H1.2 are quantified and normalized to GAPDH protein levels to indicate the release of PAR and H1.2 from nuclei after NMDA (**C**) and OGD (**E**) challenges. $P<0.05$ (E), $P<0.01$(C, H1.2), $P<0.001$(C, PAR) by one-way ANOVA, $^{***}p<0.001$ and $^{\#\#\#}p<0.001$ when compared between indicated groups by Bonferroni's posttest.

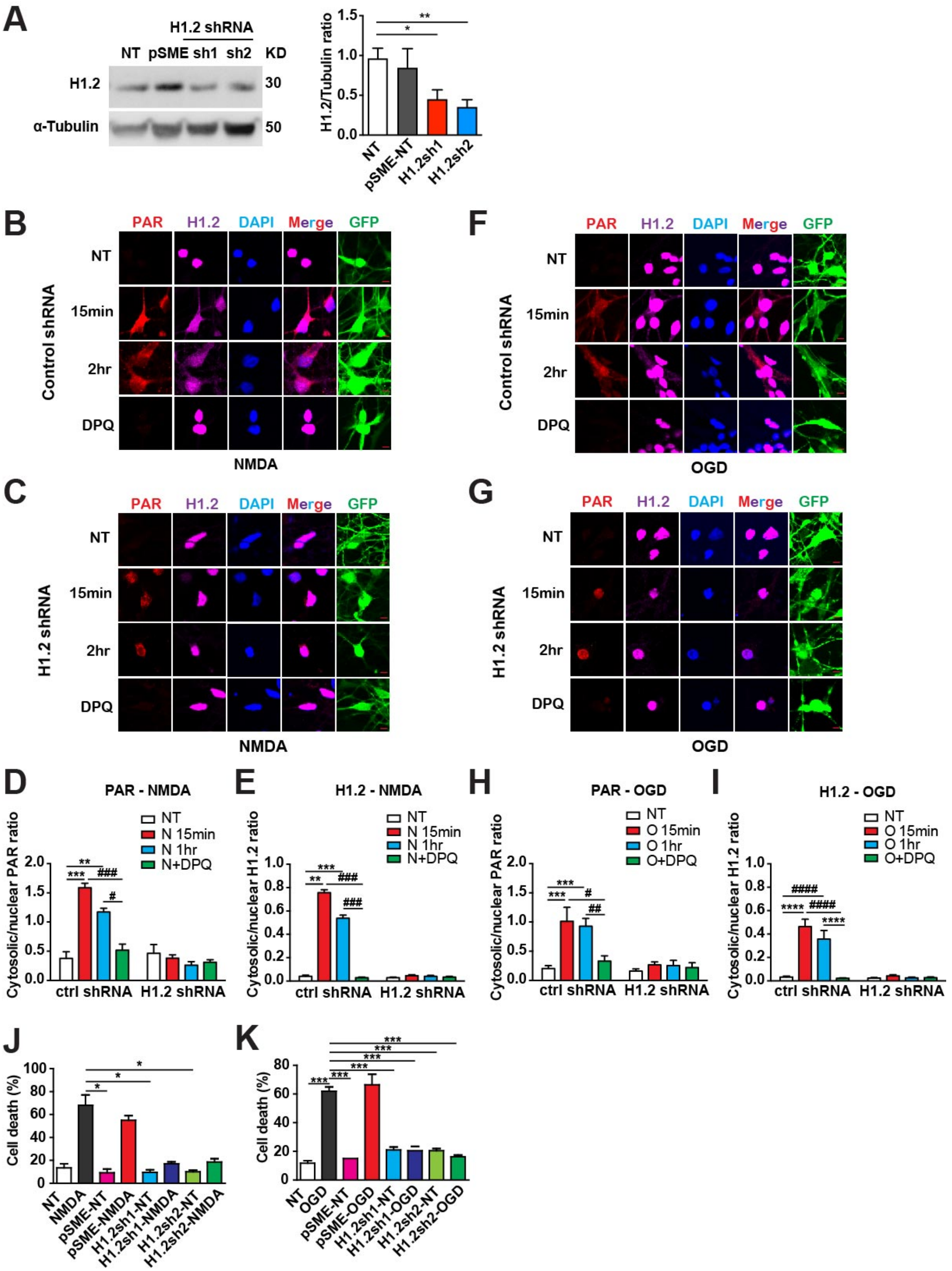


**Fig. S3, H1.2 shRNA-mediated knock down reduces PAR/H1.2 translocation and neuronal death induced by NMDA and OGD.**

(A) Representative immunoblots showing the lysates of human cortical neurons transduced with lentiviruses carrying pSME vector, and shRNAs #1 and #2 targeting H1.2. Blots were probed with anti-H1.2 antibody and alpha-Tubulin. The protein levels of H1.2 are quantified and normalized to alpha-Tubulin protein levels. The ratios are expressed as mean ± s.e.m of 4 independent experiments. **p*<0.5, ***p*<0.001, when compared to no transduction (NT) group by unpaired two-tailed t-test. ns, not significant; NT, no transduction.

(**B-E, F-I**) Representative photomicrographs showing human cortical neurons transduced with lentiviruses carrying control shRNA only (**B, F**) or with mixed shRNAs 1 and 2 targeting H1.2 (**C, G**). Cells were fixed with 4% PFA 15 minutes after NMDA (**B-E**) or OGD (**F-I**) challenges, stained with anti-PAR antibody (red), anti-H1.2 antibody (magenta) and DAPI (blue). The merged panel is a composite of the red, magenta and blue channels. The cytosolic and nuclear levels of H1.2 and PAR are quantified and the ratios (translocation) are expressed as mean ± s.e.m of 15 independent cells from 3 independent experiments. *P*<0.001 by one-way ANOVA for gene, treatment, and interaction of both PAR and H1.2 translocations in NMDA (**D**, **E**) and OGD (**H**, **I**) experiments, ***p*<0.01, ****p*<0.001, *****p*<0.0001, $^{\#}p<0.05$, $^{\#\#}p<0.001$, $^{\#\#\#}p<0.001$, $^{\#\#\#\#}p<0.001$ when compared between indicated groups by Bonferroni's posttest. NT, no treatment. Scale bar = 10 µm.

(**J, K**) Human cortical neurons are transduced with lentiviruses carrying pSME only or two shRNAs targeting H1.2, H1.2sh1 and H1.2sh2 respectively. Percentages of cell death were assessed by PI/Hoechst stain 24 hrs after NMDA (**J**) or OGD (**K**) insults, quantified and expressed as mean ± s.e.m of 4 independent experiments. *P*<0.001 by one-way

ANOVA for both NMDA and OGD experiments, *$p<0.01$, ***$p<0.001$, when compared between indicated groups by Bonferroni's posttest. NT, no treatment.

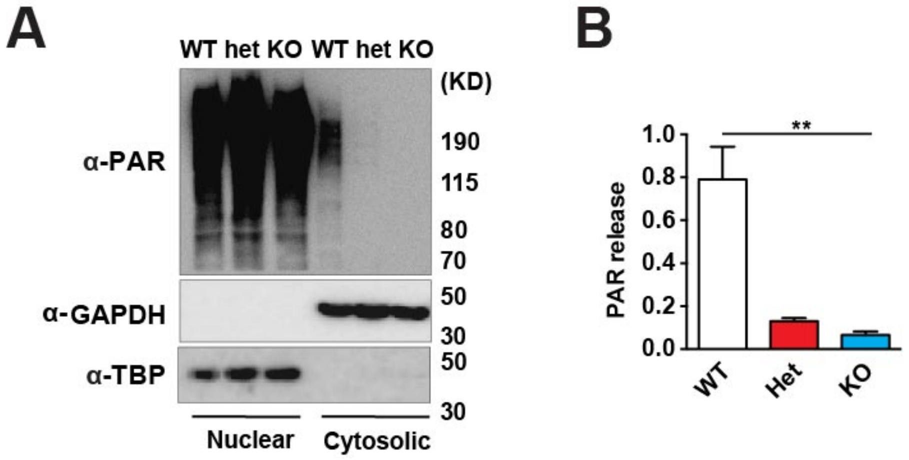


**Fig. S4. The release of PAR polymer after MNNG treatment to isolated nuclei of cortical tissues from wild type and Histone H1.2 knockout mice**

Representative immunoblots showing lysates and post nuclei buffer of nuclei treated by MNNG isolated from cortical tissues of wild type (WT), histone H1.2 knockout heterozygous (Het) and homozygous (KO) mice. Blots were probed with anti-PAR antibody, anti-TBP and anti-GAPDH. The releases of PAR from isolated nuclei after MNNG are quantified and the PAR/GAPDH ratios (PAR release) are expressed as mean ± s.e.m of 3 independent experiments. $P<0.01$ by one-way ANOVA, $^{**}p<0.01$ when compared to WT mice group, by Bonferroni's posttest.

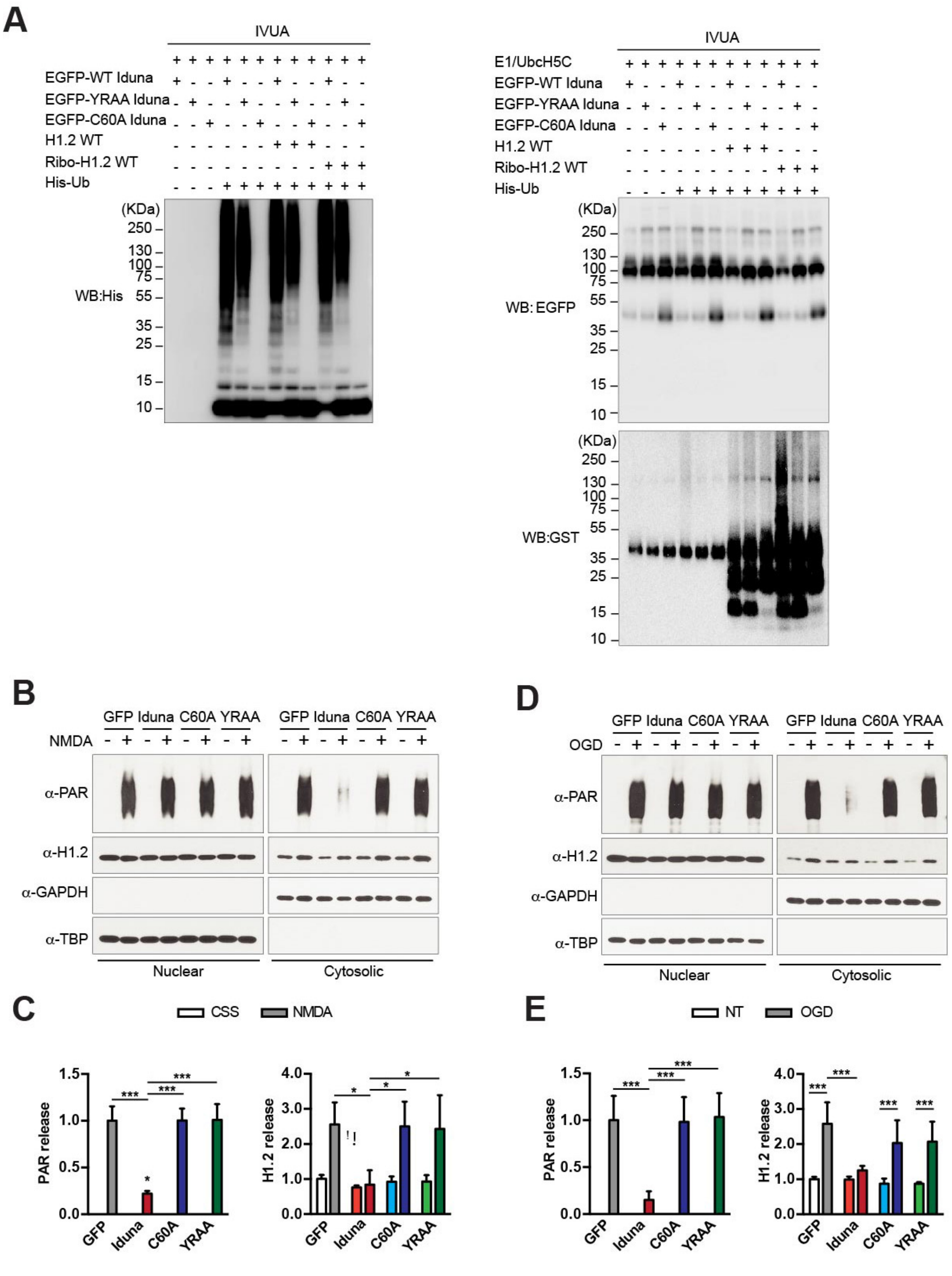


**Fig. S5. Iduna mutants representative ubiquitination blots and quantification**

**(A)** *In vitro* ubiquitination of recombinant H1.2 or ribosylated H1.2 (Ribo-H1.2) by immunoprecipitated GFP-Iduna or GFP-Iduna YRAA or GFP-Iduna C60A were analyzed by immunoblots with anti-His, anti-EGFP and anti-GST antibodies.

(**B-E**) Representative immunoblots showing nuclear and cytosolic fractions of human cortical neurons transduced with lentiviruses carrying GFP, Iduna-GFP, C60A-GFP, YRAA-GFP, and with or without NMDA (**B, C**) or OGD (**D, E**) treatments. Blots were probed with anti-PAR, anti-H1.2, anti-TBP and anti-GAPDH antibodies. The protein levels of PAR and H1.2 are quantified and normalized to GAPDH protein levels. The ratios are expressed as mean ± s.e.m of 4 independent experiments of NMDA (**C**) and OGD (**E**). $P<0.01$ by one-way ANOVA, *$p<0.05$, when compared to the control (Ctrl, Cas9 only) group by Bonferroni's posttest. NT, no transduction. The cytosolic PAR to GAPDH ratios (PAR releases) are expressed as mean ± s.e.m of 7 independent experiments. $P<0.01$ or $P< 0.001$ for genes and treatments by two-way ANOVA, ***$p<0.001$, *$p<0.05$, when compared to no treatment (NT) groups by Bonferroni's posttest.

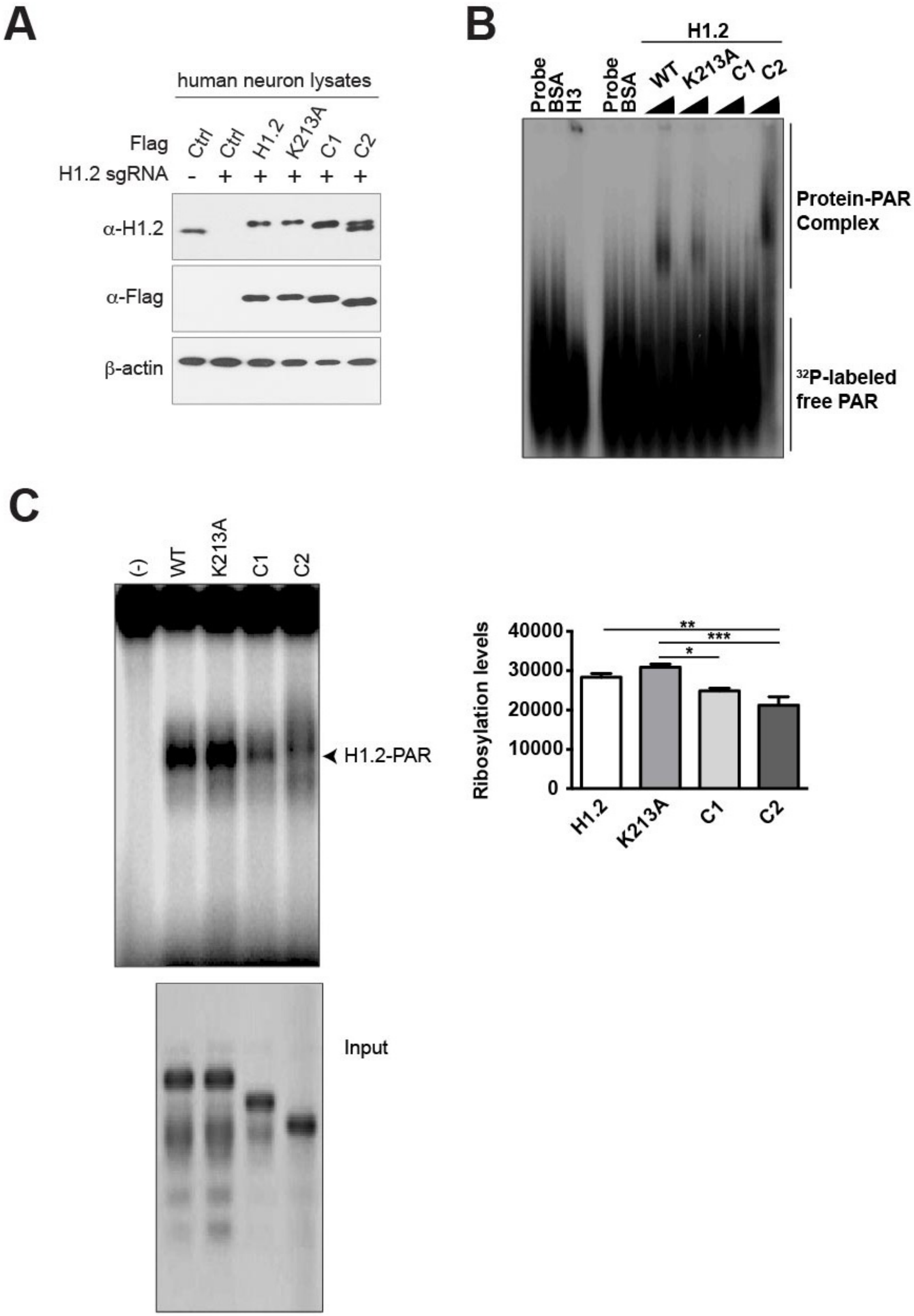


**Fig. S6. H1.2 mutants representative PAR-binding and ribosylation blots**

(A) Human cortical neurons were transduced with lentivirus carrying gRNAs targeting H1.2 followed by overexpression of Flag-tagged control or H1.2 wild type or H1.2 H213A or H1.2 C1 or H1.2 C2. Blots were probed with anti-H1.2, anti-Flag, and beta-actin antibodies.

(**B)** PAR binding capacity of H1.2-WT or H1.2-K213A or H1.2-C1 or H1.2-C2 was tested by Electromobility shift assay. Both [$^{32}$P]-PAR bound signal and unbound free [$^{32}$P]-PAR signal were separated and visualized by autoradiography. Recombinant H3 was used as a positive control and BSA was used as a negative control.

**(C**) Representative image of the *in vitro* ribosylation reaction of H1.2-WT or H1.2-K213A or H1.2-C1 or H1.2-C2 by PARP1. Each ribosylation reaction was performed in the absence of activated DNA and separated by SDS PAGE. The amount of protein used in each lane was visualized by Coomassie staining. Experiments were repeated three times and levels of ribosylation on each protein were quantified followed by autoradiography. $P<0.001$ by one-way ANOVA, *$p<0.05$, **$p<0.01$, ***$p<0.001$ when compared to H1.2 group or between indicated groups by Bonferroni's posttest. NT, no treatment.

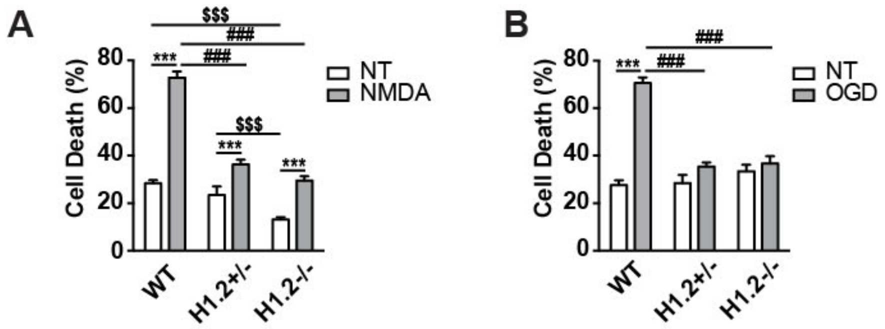


**Fig. S7. Cultured cortical neurons of Histone H1.2 knockout mice are resistant to NMDA and OGD insults**

**(A, B)** Primary cortical neuron cultures of H1.2 knockout mice are resistant to NMDA (**A**) or OGD-induced (**B**) death. Quantitative data is shown as mean ± s.e.m. of PI/Hoechst positive cell ratio by staining of three independent cultures and experiments. $P<0.0001$ by two-way ANOVA for gene, treatment, and interaction of both NMDA and OGD experiments, $^{###}$ or *** or $^{\$\$\$}p<0.001$ when compared to NT or between indicated groups by Bonferroni's posttest.
